# Substrate conformational dynamics drive structure-specific recognition of gapped DNA by DNA polymerase

**DOI:** 10.1101/263038

**Authors:** Timothy D. Craggs, Marko Sustarsic, Anne Plochowietz, Majid Mosayebi, Hendrik Kaju, Andrew Cuthbert, Johannes Hohlbein, Laura Domicevica, Philip C. Biggin, Jonathan P. K. Doye, Achillefs N. Kapanidis

## Abstract

DNA-binding proteins utilise different recognition mechanisms to locate their DNA targets. Some proteins recognise specific nucleotide sequences, while many DNA repair proteins interact with specific (often bent) DNA structures. While sequence-specific DNA binding mechanisms have been studied extensively, structure-specific mechanisms remain unclear. Here, we study structure-specific DNA recognition by examining the structure and dynamics of DNA polymerase I (Pol) substrates both alone and in Pol-DNA complexes. Using a rigid-body docking approach based on a network of 73 distance restraints collected using single-molecule FRET, we determined a novel solution structure of the singlenucleotide-gapped DNA-Pol binary complex. The structure was highly consistent with previous crystal structures with regards to the downstream primer-template DNA substrate; further, our structure showed a previously unobserved sharp bend (~120°) in the DNA substrate; we also showed that this pronounced bending of the substrate is present in living bacteria. All-atom molecular dynamics simulations and single-molecule quenching assays revealed that 4-5 nt of downstream gap-proximal DNA are unwound in the binary complex. Coarsegrained simulations on free gapped substrates reproduced our experimental FRET values with remarkable accuracy (<ΔFRET> = -0.0025 across 34 independent distances) and revealed that the one-nucleotide-gapped DNA frequently adopted highly bent conformations similar to those in the Pol-bound state (ΔG < 4 kT); such conformations were much less accessible to nicked (> 7 kT) or duplex (>> 10 kT) DNA. Our results suggest a mechanism by which Pol and other structure-specific DNA-binding proteins locate their DNA targets through sensing of the conformational dynamics of DNA substrates.

**Significance Statement:** Most genetic processes, including DNA replication, repair and transcription, rely on DNA-binding proteins locating specific sites on DNA; some sites contain a specific sequence, whereas others present a specific structure. While sequence-specific recognition has a clear physical basis, structure-specific recognition mechanisms remain obscure. Here, we use single-molecule FRET and computer simulations to show that the conformational dynamics of an important repair intermediate (1nt-gapped DNA) act as central recognition signals for structure-specific binding by DNA polymerase I (Pol). Our conclusion is strongly supported by a novel solution structure of the Pol-DNA complex wherein the gapped-DNA is significantly bent. Our iterative approach combining precise single-molecule measurements with molecular modelling is general and can elucidate the structure and dynamics for many large biomachines.

---

Protein machines functioning on chromosomes and plasmids utilise different mechanisms to locate their targets on DNA. Sequence-specific DNA-binding proteins, such as restriction enzymes and transcription factors, recognize a particular nucleotide sequence via a combination of direct and indirect readouts (1). Whereas direct readout involves specific interactions between the DNA bases and protein amino acid side chains, indirect readout senses sequence-dependent structural and mechanical features, such as major or minor groove width, and conformational flexibility (2). In contrast, structure-specific proteins have no sequence specificity; instead, they interact with particular DNA structures (e.g., gapped duplexes, and 5’ or 3’ overhangs). While sequence-specific DNA binding mechanisms have been studied extensively, structure-specific mechanisms remain unclear.

Many enzymes involved in DNA repair and replication are necessarily structure-specific, and have been shown to interact with bent DNA substrates; examples include Flap endonuclease 1 (3-5), DNA Polymerase β (6), XPF (7), and MutS (8, 9). Although catalytic reasons for substrate distortion have been suggested for individual systems, it is unclear whether the structure and dynamics of bent states serve as general recognition signals for binding or substrate selectivity. A key related question is whether these proteins induce DNA bending upon binding (via an “induced” fit mechanism) or they recognise a pre-bent state adopted by the DNA prior to protein binding (a “conformational selection” mechanism), or a combination of both.

Here we studied the *E. coli* DNA polymerase I (Klenow Fragment; Pol), a structure-specific protein responsible for Okazaki fragment processing in lagging-strand DNA replication, as well as for DNA synthesis during DNA repair. In both roles, the polymerase recognizes and binds to a gapped DNA substrate and polymerizes across the gap. After gap filling, strand-displacement synthesis may follow; this is important for Okazaki fragment processing, as the polymerase continues to synthesize DNA whilst displacing an RNA primer, which is subsequently excised.

Attempts to understand the DNA binding and recognition mechanism for Pol are complicated by the absence of crystal structures of DNA-Pol binary complexes containing downstream duplex DNA, by the heterogeneity of Pol-DNA complexes (10, 11), and by the conformational mobility of the free DNA substrate. As a result, there are many open questions regarding the mechanisms of strand-displacement DNA synthesis and substrate recognition.

We investigated the mechanism of structure-specific recognition by Pol via a combination of single-molecule Förster resonance energy transfer (smFRET) and molecular modelling. The single-molecule nature of our work addressed the issues of conformational and compositional heterogeneity, and led to a FRET-restrained solution structure of the binary complex, Pol bound to 1-nt gapped DNA. This structure revealed a substantial bend in the DNA substrate (which was also supported by complementary FRET experiments in living bacteria), and provided insight into protein and DNA structural features crucial for strand-displacement synthesis. The structure also served as the starting point for atomistic molecular dynamics simulations, which revealed the dynamic nature of the binary complex and the specific interactions between the protein and DNA. Experimental smFRET measurements and coarse-grained modelling allowed us to characterise the conformational ensemble of the free substrate and propose a mechanism for substrate recognition and binding by DNA polymerase I, which is likely to apply to many other structure-specific DNA-binding proteins.

## Results

### Structural analysis of multiple species in dynamic equilibrium

To analyse the structure of the binary complex of Pol with a 1-nt gapped DNA in solution, we determined numerous DNA-DNA and protein-DNA distance restraints within freely diffusing Pol-DNA complexes using single-molecule confocal fluorescence microscopy combined with alternating-laser excitation (12-14). We measured DNA-DNA distances between labelled sites in the upstream and downstream duplex regions of gapped DNA containing a 3’-dideoxy nucleotide (to prevent any chemistry occurring; Fig 1A). We also measured DNA-protein distances between a FRET donor dye attached to one of three Pol residues (K550C, L744C, C907; Fig 1B) and a FRET acceptor dye attached to one of 13 labelling sites on gapped DNA. Pol activities were not significantly affected by the dye presence (10, 15).

**Figure 1.**
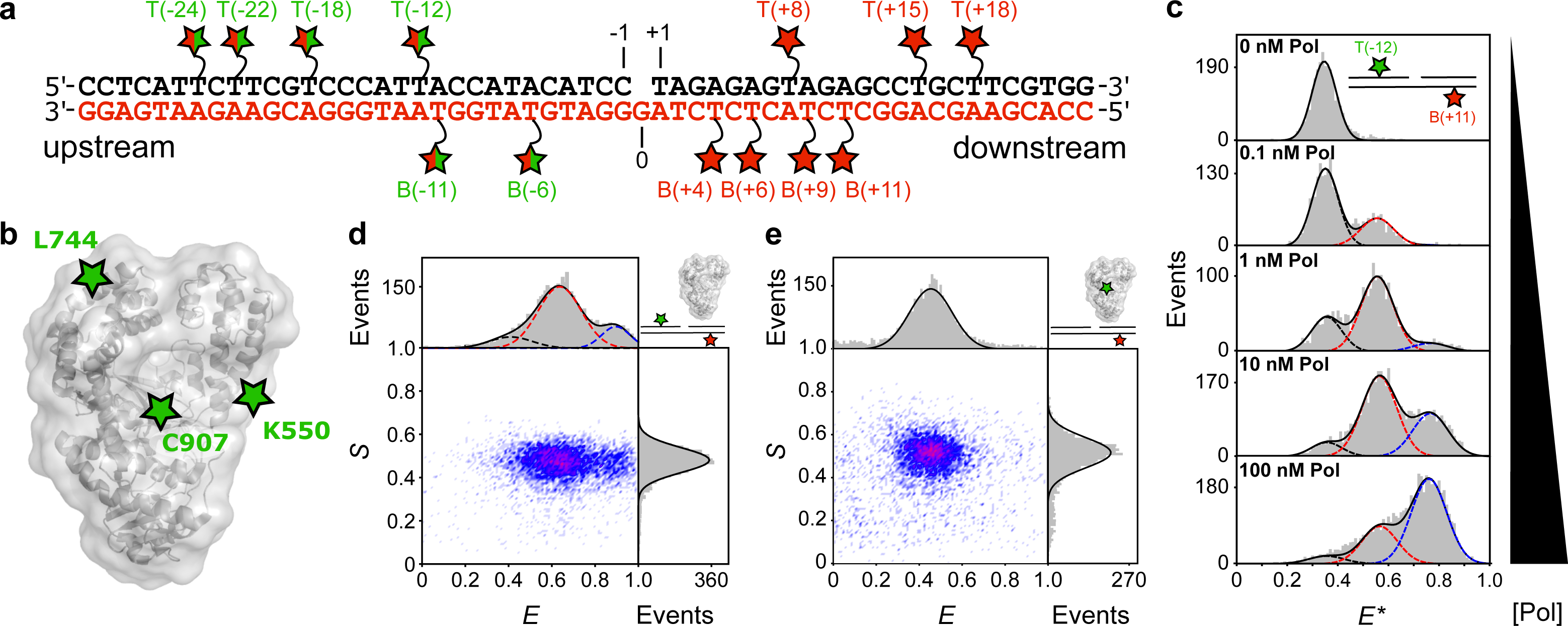
Measuring distances within single polymerase-DNA complexes in heterogeneous mixtures with dynamic species. (a) Schematic of a 1-nt gapped DNA substrate showing the template (red lettering) and non-template strands (black). Red stars represent acceptor labelled dT bases. Split red/green stars indicate positions labelled with donor for DNA-DNA FRET, or acceptor for DNA-protein FRET. (b) DNA Polymerase I (Klenow Fragment; Pol) structural schematic (grey - pdb 1KLN) and donor labelling positions (green stars) (c) Apparent FRET histograms for the doubly-labelled substrate T(−12) B(+11) at increasing concentrations of Pol. The data (grey bars) were fitted with up to three Gaussians (black, red and blue dashed lines), yielding apparent FRET efficiencies, E* of 0.35, 0.55 and 0.75. (d) Corrected ES histogram for a DNA-DNA FRET measurement (here, for T(−12)B(+11) in the presence of 3 nM Pol). Data (grey bars) were fitted by the sum of three Gaussians (solid black lines) centered on E=0.41 (black dash), E=0.63 (red dash) and E=0.90 (blue dash) respectively. (e) Corrected ES histogram for a protein-DNA FRET measurement (here, for C907-Cy3B B6-Atto647N). Data were fitted with a single Gaussian function, centered on E=0.48. See also Fig S1.

We determined the Pol concentration required to form binary complexes by monitoring FRET between upstream and downstream sites on the DNA at increasing Pol concentrations. Three distinct FRET states were observed during the titration, indicating three different conformations of the gapped DNA (Fig 1C). A low-FRET state corresponding to free DNA; a high-FRET state corresponding to a (Pol)2-DNA ternary complex; and a mid-FRET state corresponding to two distinct species, the Pol-DNA binary complex, and a (Pol)_2_-DNA ternary complex, with indistinguishable FRET signals (see *Methods*; Fig S1A-B). The increased FRET for the Pol-DNA binary complex vs. free DNA suggested substantial DNA bending in the binary complex. Using global fitting *(Methods)*, we recovered dissociation constants of *K*_D1_ = 360 ± 60 pM for the binary complex, and *K*_D2_ = 9 ± 4 nM for formation of the mid-FRET (Pol)_2_-DNA dimer species (Fig S1B).

To characterize the structure of the DNA in the binary complex, we set the Pol concentration to 1 nM thereby maximising the population of the binary complex (Fig 1D), and measured 34 DNA-DNA FRET distances (Table S1, and Fig S1). We also obtained 39 protein-DNA distance restraints for the binary complex, measuring FRET between donor-labelled Pol and acceptor-labelled DNA (Table S1 and Fig 1E). Single-molecule confocal measurements are generally limited to a maximum concentration of fluorescent species of ~100 pM, however, because of its low dissociation constant, a detectable quantity of the binary complex was present even at 100 pM dye-labelled Pol (Fig 1E).

### The DNA substrate is bent by 120° in the Pol-DNA binary structure

To obtain structural models of the Pol-DNA complex, we used our 73 distance restraints to perform rigid-body docking using Pol and two shortened DNA helices representing the upstream and downstream DNA (Fig 2A). We generated 32 refined structures and ranked them according to their fit to the measured distances (see *Methods).* A single model (Fig 2B) emerged with a significantly better fit than the other structures (Fig S2A). In this model, the position of the upstream DNA agreed very well with the position of a DNA fragment in a crystal structure of a Pol-DNA binary complex (16), (RMSD = 2.9 Å; Fig S2B), demonstrating the accuracy of our structural model. To test the robustness of our model, we generated 100 ‘bootstrapped’ structures by randomly perturbing the 73 distance restraints in proportion to their experimental errors, repeating the docking calculations, and calculating the RMSD for each DNA backbone phosphate atom across all bootstrapped structures (average RMSD = 3.8 Å, Fig S2E).

**Figure 2.**
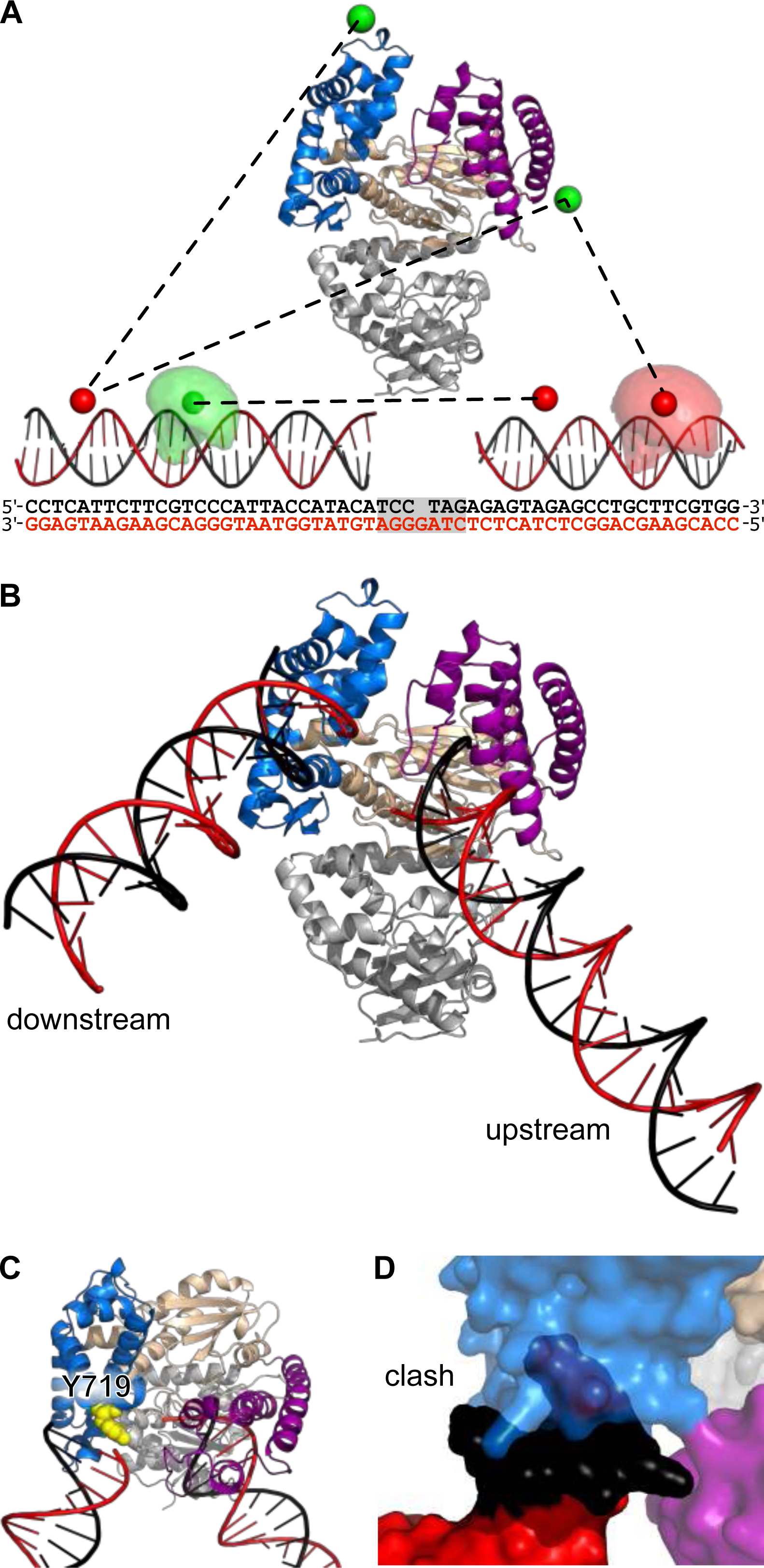
Pol-DNA binary structure from rigid-body docking. (a) Pol structure showing the fingers (blue), thumb (purple) and palm (wheat) subdomains and the proof-reading exonuclease domain (grey). Example DNA-DNA and protein DNA-distances (black dashed lines) are shown between mean dye positions (green and red spheres). Example accessible volumes of a donor (pale green cloud) and an acceptor (pale red cloud) dye are also shown, along with the full sequences of the docked DNAs; the shaded region indicating the DNA not used for the docking. (b) Results of the rigid-body docking: template DNA (red), non-template DNA (black), subdomains coloured as in (A). (c) Position of Y719 relative to downstream DNA. (d) Clash between full-length downstream DNA and the fingers subdomain (cyan). See also Fig S2 and Table S1.

Having established the accuracy and precision of our model, we inspected it for insights into the DNA-binding and strand-displacement mechanisms. The most striking feature was the significant kink in the DNA substrate, a ~120° bend compared to straight duplex (Fig 2B). Further, the downstream DNA is positioned close to the fingers subdomain (Fig 2C), with the helical axis aligned with Y719 (*Bst* numbering used as default; this corresponds to residue F771 in *E. coli).* Substitution of this residue with alanine was previously shown to significantly impair strand displacement synthesis by Pol (17). Extending the downstream DNA to its full length by modelling the previously deleted base pairs proximal to the gap, resulted in a clash between the additional DNA and the Pol fingers (Fig 2D) indicating that the gap-proximal downstream DNA may be partially melted in the binary complex.

We also used 21 FRET restraints (Table S1) to obtain the relative orientation of the upstream and downstream DNA in the high-FRET (Pol)2-DNA ternary complex and generated a low-resolution model for this complex in which the DNA was more severely bent (by ~140°; Fig S2F; RMSD = 12 Å, Fig S2G).

### All-atom MD simulations give structural insights into the strand displacement mechanism

To probe the exact position of DNA in the binary complex, its dynamics and any specific contacts with Pol, we carried out all-atom MD simulations. We generated five different starting models by combining the DNA from our FRET-restrained structure with the short DNA fragment present in the 4BDP Bst X-ray structure (18) (see *Methods* and Fig S3B), and performed two unconstrained 100-ns simulations from each model.

Whereas the DNA fragment present in the X-ray structure remained stably bound to Pol (RMSD 2.8 ± 0.8 Å), the upstream and downstream segments flanking this DNA were much more mobile (RMSD 11.1 ± 4.4 Å and 19.2 ± 8.8 Å, respectively; Fig S3C), with the end-to-end DNA distance ranging from 24 to 144 Å (Fig 3A and Fig S3D). The first six nucleotides of the downstream, nontemplate DNA (nucleotides T(+1) to T(+6), termed the non-template flap) also displayed appreciable dynamics (RMSD 8.7 ± 3.7 Å, Fig 3B and Fig S3E), and did not dock in a particular conformation. To study the extent of DNA melting in this region, we counted the hydrogen bonds formed between the six nucleotides of the non-template flap and the template strand: for most of the simulation time, 2 or 5 hydrogen bonds were present (Fig 3B), corresponding either to a single AT or to an A-T plus a G-C pair, respectively, consistent with base-pairing of the two nucleotides at the base of the flap. Hence, in most conformations, 4 or 5 nucleotides of the flap were melted. Contacts between the flap and Pol were transient and diverse in terms of the residues involved; the most consistent interactions were sequence-unspecific, being formed between DNA phosphates and positively charged residues (mainly R729 and K730).

**Figure 3.**
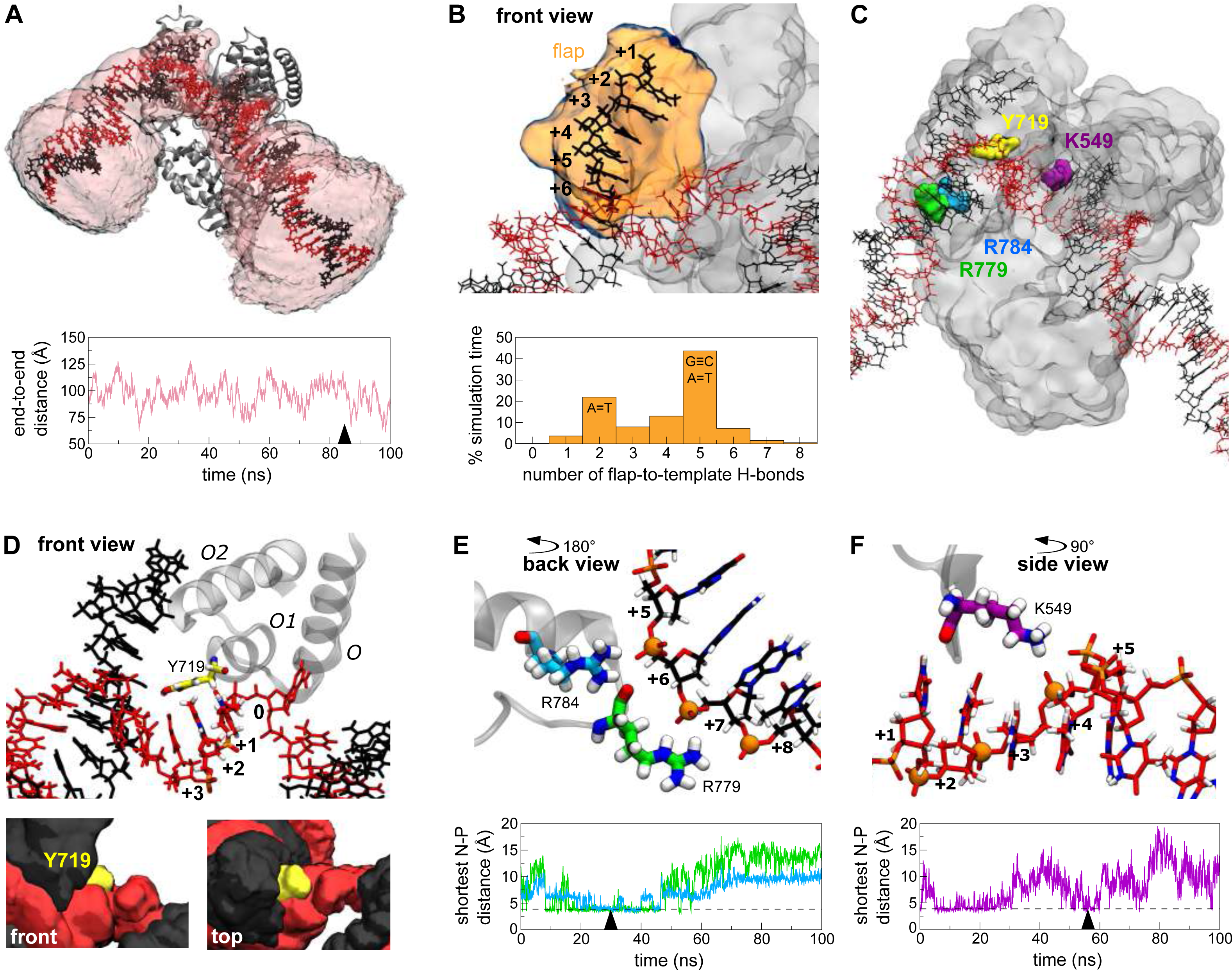
Binary complex structure and dynamics. (a) Representative snapshot of the DNA-Pol binary complex from a 100-ns MD simulation, showing the volume accessed by the DNA over the simulation (pale pink). The plot shows the DNA end-to-end distance fluctuations over the same simulation, with the ends taken as the terminal non-hydrogen atoms of the template strand. The time point corresponding to the snapshot is indicated with an arrowhead. (b) Representative snapshot of the conformation of the 6-nt non-template flap, with its volumetric map during a 100-ns simulation (pale orange). The plot shows the frequency of the number of hydrogen bonds formed between the flap and the template strand of downstream DNA during the entire 1-μs (10× 100 ns) simulation time. (c) Overview of Pol residues involved in strand separation or interactions with downstream DNA. See also panels (D) to (F). (d) Involvement of Y719 in strand separation of downstream DNA. Top - a representative snapshot of the position of Y719 relative to the template DNA strand. The three DNA residues positioned closest to Y719 during the time course of the simulation are highlighted in CPK colouring. The position of the three-helix bundle is shown for reference; the rest of the protein is omitted for clarity. Lower panels, two different views of the volumetric maps of Y719 (yellow), template (red) and non-template DNA strands (black) during a 100-ns simulation. (e) A representative snapshot of the interactions between R779 (green) and R784 (cyan) with phosphate groups (orange spheres) in the non-template strand of downstream DNA. The plot shows the minimum distance between the side-chain nitrogen atoms of R779 (green) or R784 (cyan) to any phosphorous atom in the non-template strand of downstream DNA during a 100 ns simulation. Arrowhead denotes the time point corresponding to the snapshot, and dashed lines indicate the distance corresponding to an interaction. (f) A representative snapshot of the interaction between residue K549 (purple) with phosphate groups in the template strand of downstream DNA. See also Fig S3, Movie M1 and PBD File P1.

Many of the protein-DNA interactions in the active site (Y714, S717, Y719 and R789, all contacting the template strand) were similar to those observed in the X-ray structure (18). Further, in our simulations, the conserved three-helix bundle (O, O1 and O2 helices in the fingers subdomain), and especially residue Y719 (F771 in *E. coli*) were consistently positioned between the downstream template and non-template strands (Fig 3D); Y719 was typically positioned perpendicular to bases B(+1) and B(+2) of the template strand (Fig 3D, upper panel), and occasionally stacked against them (Fig S3F). The position of Y719 is consistent with a previously suggested mechanism in which Y719 acts as a “wedge”, separating the non-template strand from its template counterpart (17). Despite the intrinsic dynamics of the non-template strand, the stable positioning of Y719 against the template strand likely prevents re-pairing during catalysis.

Finally, we observed interactions between downstream DNA and the polymerase, which consistently involved positively charged residues on the Pol surface and the negatively charged phosphate groups of the DNA backbone. These interactions occurred in two regions: the first involved R779 (S831) and R784 (R836) that contact the duplex region of downstream DNA (Fig 3E), and the second featured K549 (K601) of the thumb region interacting with the unpaired template strand (Fig 3F). Whilst any individual nitrogen-phosphate interaction was transient, each residue contacted up to 6 phosphate groups, resulting in Pol-downstream DNA interactions persisting for most of the simulation time. The dynamic nature of these interactions likely reflects the need for rapid Pol movement along its DNA substrate during DNA synthesis.

**Downstream DNA is melted in the DNA-Pol binary complex.** To study the melting of the downstream non-template strand predicted by both our docked binary complex model (Fig 2D) and MD simulations (Fig 3B), we used quenchable FRET (quFRET), a single-molecule assay able to detect local DNA unwinding (19-21). In quFRET, when the donor (Cy3B) and acceptor (Atto647N) are in close proximity (< 2 nm), their emission is quenched, yielding only few events with intermediate stoichiometry (0.4 < S < 0.8) (see *Methods)*. Upon local DNA melting, the two dyes move further apart and the quenching is reduced, leading to a large increase in both the number, and proportion of events with intermediate stoichiometry (mostly occurring at high FRET efficiencies, as the inter-dye distance remains short).

We studied a 1-nt gapped DNA substrate labelled with donor and acceptor dyes at positions T(+1) and B(+4), respectively. In the absence of Pol, the dyes are in very close proximity; as a result, we detected few intermediate-S events (Fig 4), comprising only ~25% of all acceptor-containing molecules (Fig S4A). On addition of Pol, we observed a ~4.5-fold increase in the number of such events per measurement, with a peak at high FRET (E* > 0.9; Fig 4), now comprising ~75% of all acceptor-containing molecules (Fig S4B). These results demonstrate an increase in dye separation and reduced quenching, consistent with the presence of local melting at the 5’-end of the downstream non-template strand in the binary complex.

**Figure 4.**
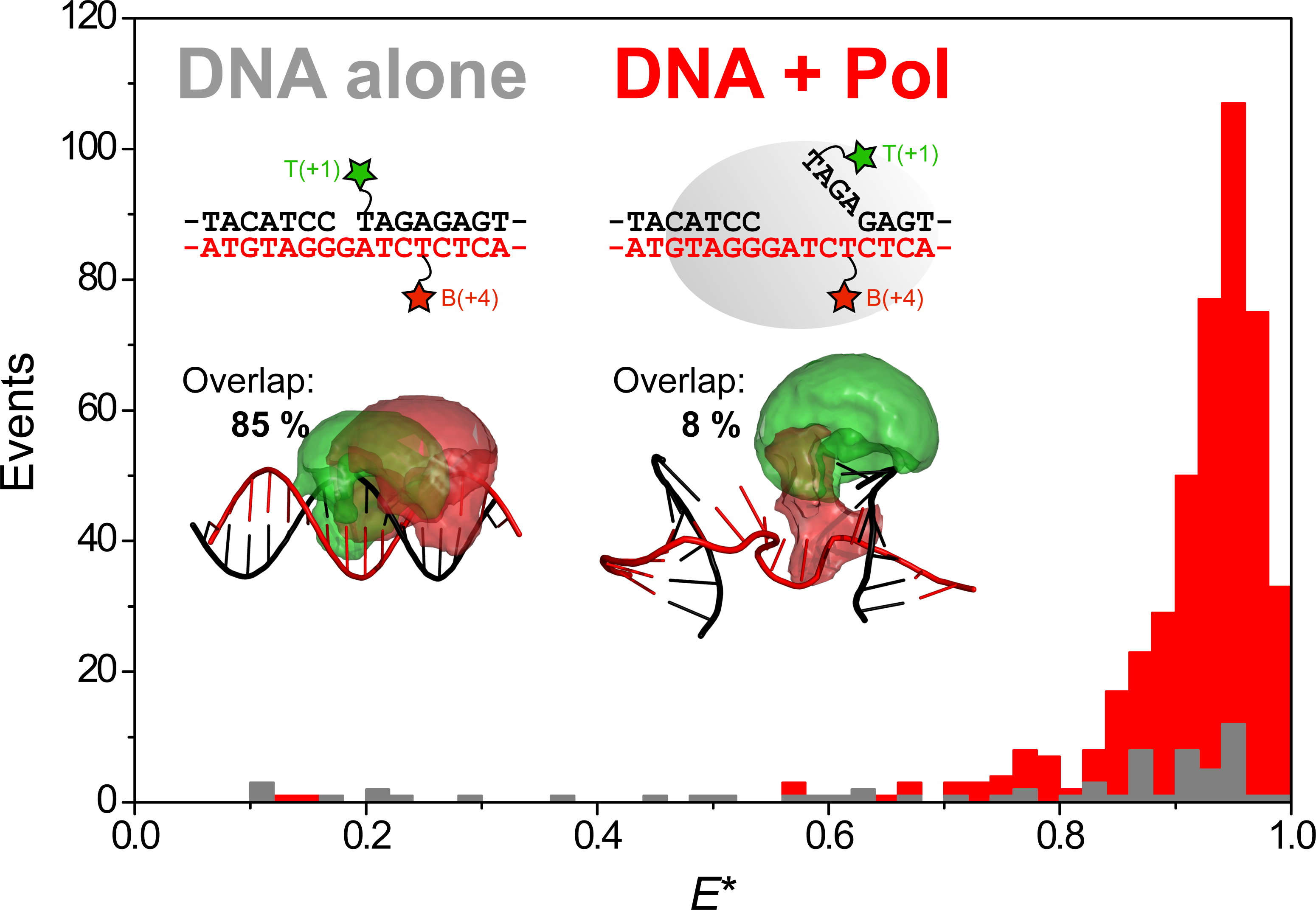
Downstream DNA is melted in the binary complex. Results from quenchable FRET experiments. The plot shows the number of events with mid-stoichiometry (0.4 < S < 0.8) vs the apparent FRET efficiency, for DNA substrate T(+1)B(+4) alone (grey bars) and in the presence of 3 nM Pol (red bars). Inset: Schematics of the labeling positions and DNA structures for the unbound (left; B-DNA) and bound conformations (right; snapshot from MD simulations, atomic coordinates provided as SI), and the related accessible volumes of the donor (green) and acceptor (red) dyes, quoting the percentage overlap between them (see main text). See also Fig S4.

To monitor the extent of melting along the downstream DNA, we tested a substrate with donor and acceptor dyes at B(+9) and T(+8) respectively. For these labelling positions, we observed similar quenching in both the absence and presence of Pol (Fig S4D-F), implying that this DNA site remains base-paired in the binary complex. This suggests the maximum number of melted base-pairs is in the binary complex is seven, consistent with the 4-5 observed in our MD simulations.

To relate the observed quenching changes to DNA rearrangements upon Pol binding, we analysed the accessible volumes (AVs) of the dyes in both bound and unbound conformations. Assuming a B-DNA duplex conformation for the unbound substrate, 85% of the donor AV overlapped with the acceptor AV (Fig 4 - inset); this was reduced to 8% in representative MD snapshots of the binary complex in which 4-5 nt are melted. Additionally, no quenching was observed for substrates with dye labelling positions with 0% AV overlap (Fig S4G-I). Hence, the observed reduction of quenching in the bound state correlated well with the change in AV overlap. Taken together, our quFRET results strongly support the hypothesis that the downstream duplex DNA is partially unpaired in the binary complex.

**The free 1-nt gapped DNA substrate adopts bent and frayed states**. To examine to what extent DNA bending and downstream melting were present in the free substrate, and to establish whether Pol recognizes such structural features via conformational selection, or induces them upon binding, we studied the structure and dynamics of the 1-nt gapped DNA substrate in the absence of Pol.

We used smFRET to collect 34 DNA-DNA distances within the free DNA (Fig 5A-B and Table S2). Corrected ‘ES’ histograms showed a single FRET peak, consistent with either a single DNA conformation or rapid (sub-millisecond) conformational averaging (Fig S5A-C). Rigid-body docking of the two duplex portions of the substrate, failed to produce a unique structural model; instead, five structures of the gapped-DNA emerged, with angles between the duplex arms spanning from 8° to 25° (Fig S5D). This approach assumed a single static structure was responsible for the experimental FRET distances; however, the substrate is expected to be highly dynamic (22, 23). To take into account these dynamics, we conducted coarse-grained molecular modelling on the gapped-DNA substrate using the oxDNA model, which allows rapid and efficient conformational sampling, and has been shown to describe well the structural, thermodynamic and dynamic properties of many DNA systems (Fig S5E) (24-27).

**Figure 5.**
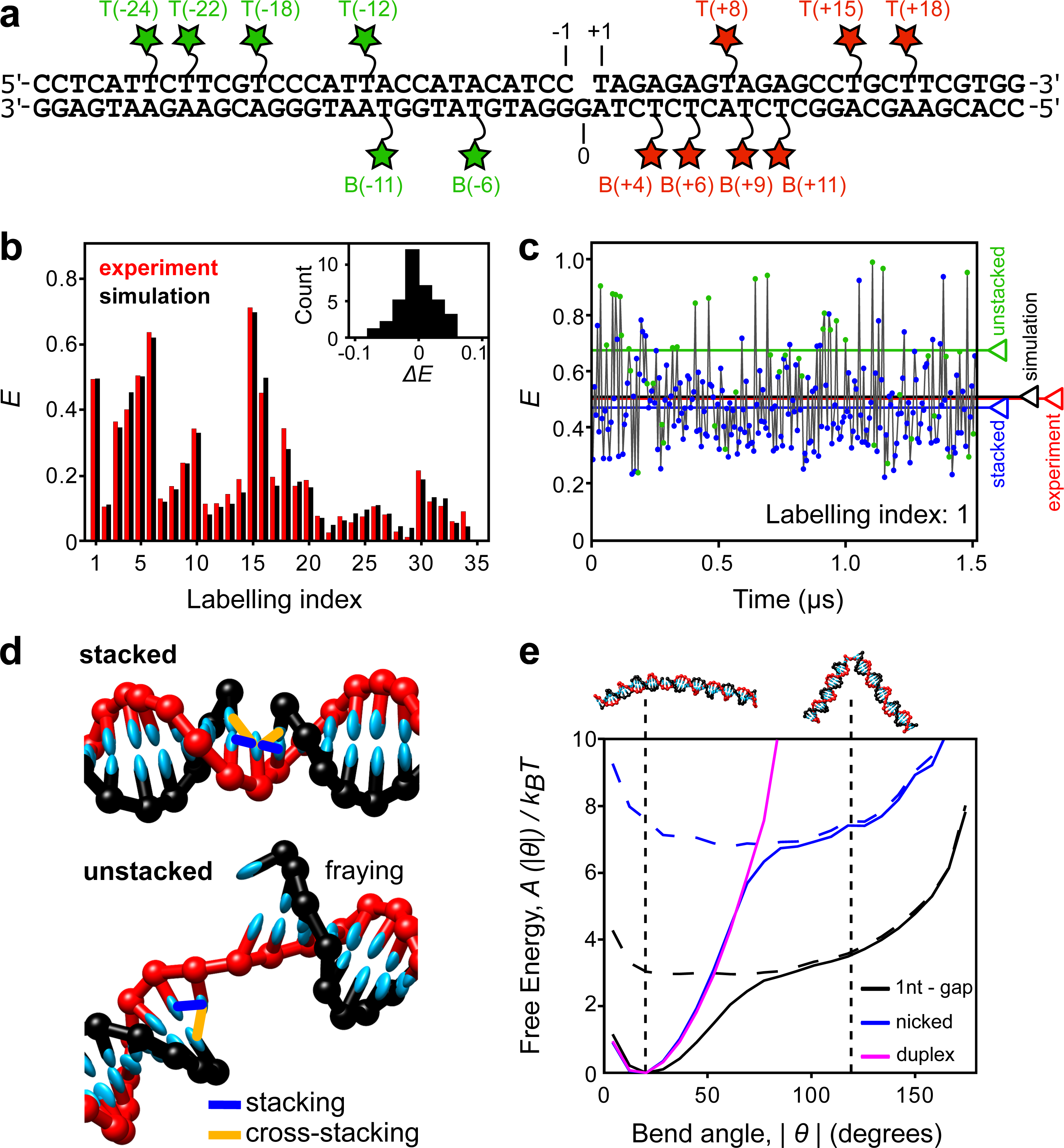
Gapped DNA substrate adopts highly bent conformations due to base unstacking. (a) Schematic of the 1-nt gapped DNA substrate. Green and red stars indicate donor and acceptor labelled dT bases, respectively. (b) Comparison of experimental (red) and simulated (black) corrected FRET values for 34 FRET measurements in the 1-nt gapped substrate (Labelling index listed in Table S2). Inset: residual histogram, ΔE = *E*_model_ - *E*_measured_ (c) Simulated FRET time trace for substrate T(+8)B(−11), labelling index 2. Stacked (blue circles) and unstacked (green circles) states are identified. The average FRET efficiency across the entire simulation is shown for each state individually (blue and green lines), for the sum of both states (black line) and for the experimentally determined FRET value (red line). (d) Typical snapshots of the stacked and unstacked states indicating the stacking (blue) and cross-stacking (orange) interactions present. (e) Free-energy profiles as a function of the bend angle (θ = 0 corresponds to a straight duplex) for 1nt-gap (black), nicked (blue) and duplex (magenta) DNAs. The total free energy (solid lines) and the contribution due to unstacked states alone (broken lines) are shown. The vertical dotted lines and cartoons correspond to the most stable bend angle (20°) and the bend angle observed in the binary complex structure (120°). See also Fig S5, Table S2 and Movie M2.

Using an adapted AV approach (see *Methods* and Fig S5G) (28, 29), we calculated the FRET efficiency arising from each dye pair at regular simulation intervals (Fig 5C). Transitions between different configurations were rapid (submicrosecond) and much faster than the temporal resolution of the smFRET experiments (~1 ms; see *Methods)*. Therefore, the average FRET efficiencies from the simulations are expected to agree with those measured experimentally. Indeed, we see excellent agreement between the experimental and modelled FRET efficiencies across all 34 measured FRET pairs (RMSD = 0.026, <ΔFRET> = -0.004; Fig 5B and Table S2, c.f. the estimated experimental error, FRET_error_ = ± 0.025, see *Methods)*. The fit to the experimental data was significantly worse for the best of the five static structures obtained from rigid-body docking (RMSD = 0.054), suggesting that the coarse-grained simulations better describe the experimental conformational ensemble and dynamics for these highly dynamic substrates.

The simulations of the free substrate identified two classes of structures: in ~80% of configurations, both stacking interactions between the three nucleotides opposite the gap were maintained, resulting in a straighter geometry (Fig 5D, top). In ~20% of configurations, at least one of these stacking interactions was broken, resulting in a state in which the system can explore a wide variety of bend angles (Fig 5D - bottom). FRET efficiencies were typically larger for unstacked configurations (Fig 5C - green circles), as the enhanced bending allowed the dyes to explore more proximal positions. Notably, although the stacked conformations are the dominant contributor to the FRET signal, excluding the unstacked states from the average FRET calculation lead to average computed FRET values significantly lower than those observed experimentally, worsening the agreement to the experiment (ΔFRET> = -0.004 with unstacked states vs -0.026 without unstacked states, Table S2). This finding strongly suggests that bent states are present in the experimental ensemble for the gapped substrate. Bent states were also detected in all-atom MD simulations on the gapped substrate (Fig S5H).

Encouraged by the excellent agreement between the computational and experimental results for the free substrate, we calculated free-energy landscapes from the relative abundance of conformations with specific bend angles (Fig 5E). The landscapes show that the angle seen in the Pol-bound state of the gapped substrate (~120°) is accessible to the unbound gapped substrate (free energy difference of <4 kT); this bend angle is achievable only upon breaking at least one of the stacking interactions. However, once the stacking is broken, the substrate can freely explore a relatively flat landscape (Fig 5E - dashed lines), where the gap acts as a hinge. Simulations on nicked and duplex DNAs showed that it is harder for these substrates to adopt bend angles of 120° (Fig 5E), due to the increased energetic cost of breaking an additional stacking interaction and, in the case of the duplex, the extra chain connectivity constraints (>7 kT - nick, >>10 kT - duplex).

We also inspected the coarse-grained simulations for evidence of melting of the downstream duplex DNA in the gapped substrate alone. In 28% of all configurations, we observed melting of the A-T base pair (fraying) at the downstream site immediately adjacent to the gap (Fig S5F); when looking only at unstacked configurations, this fraction increased to 35%, partly due to the loss of a stabilising cross-stacking interaction (Fig 5D). The second nucleotide was frayed in ~5% of configurations irrespective of stacking state (Fig S5F); fraying of three nucleotides was never observed. Taken together, our results are consistent with a conformational selection model in which Pol initially interacts with an unstacked, bent configuration of gapped DNA, a significant proportion of which is frayed by 1-2 nt around the gap.

**Bent DNA detected in live cells**. To test for the existence of bent gapped DNA *in vivo*, as suggested by our binary structure, we measured the FRET efficiencies of individual labelled-DNA substrates in live *E. coli* cells. A small number of 1-nt gapped DNA molecules (1-5 molecules per cell) were internalized into cells by electroporation (30-32) and their FRET efficiencies monitored (Fig 6A; *Methods*).

**Figure 6.**
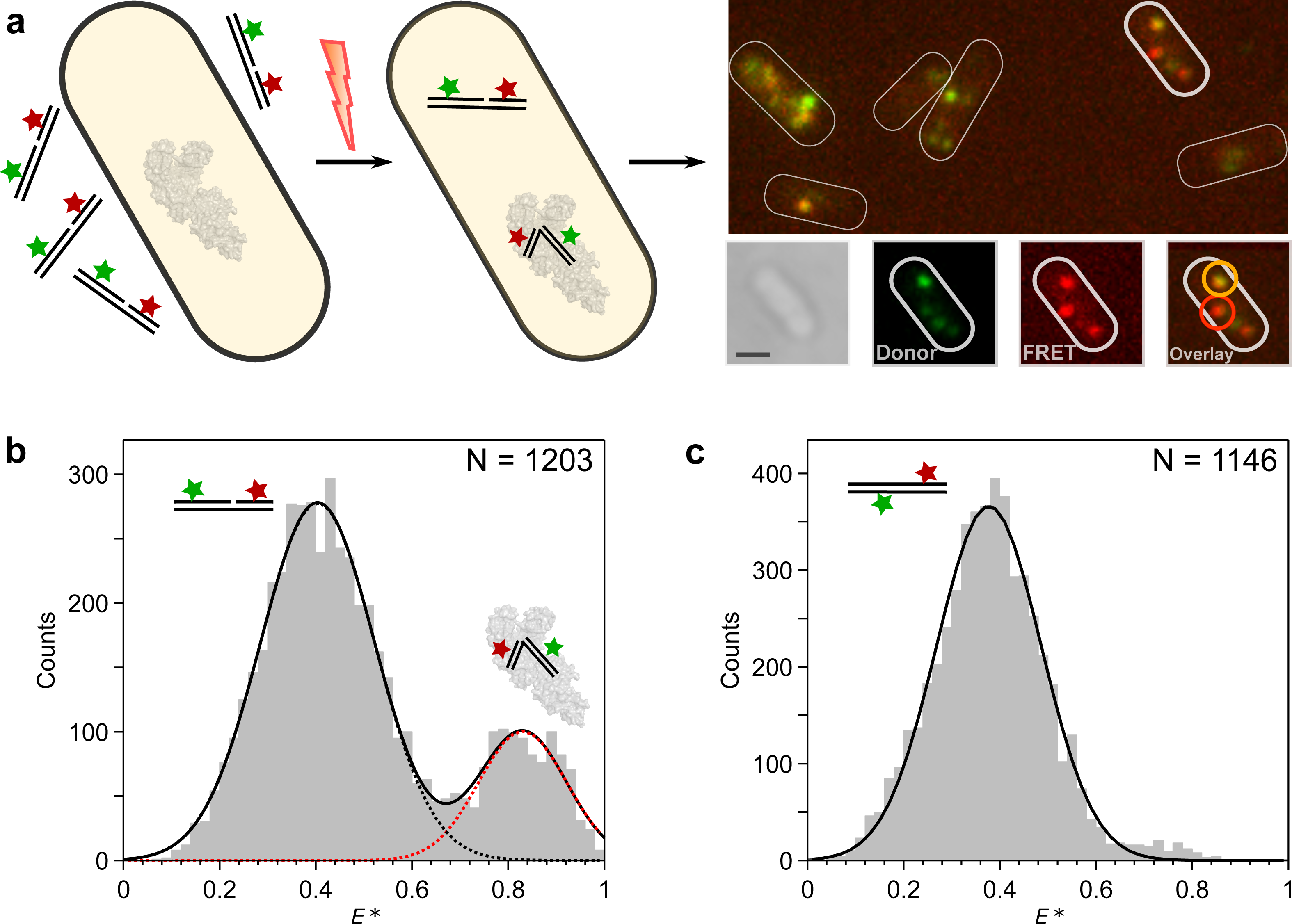
DNA bending detected in live cells. (a) Schematic showing internalization of doubly-labelled gapped DNA fragments into live *E. coli* using electroporation and single-molecule imaging (left to right). Example cell (bottom right) is shown in white-light image, donor fluorescence channel, FRET fluorescence channel, and latter both combined in overlay image. The overlay image is color-coded such that intermediate-FRET molecules appear orange and high-FRET molecules appear red; two example molecules are highlighted accordingly. Scale bar: 1 μm. (b) FRET histogram of tracked gapped DNA trajectories *in vivo*. Two major FRET species were observed for the T(−12)T(+8) substrate, which were attributed to unbent DNA (orange; *E\** = 0.40) and bent DNA (red; *E\** = 0.83). The number of trajectories (N) is stated for each experiment. (c) FRET histogram of tracked duplex DNA trajectories *in vivo*. A single low-FRET species, was observed (black, *E\** = 0.38). See also Fig S6.

We first internalized the T(−12)T(+8) gapped substrate, as it showed a large FRET change upon Pol binding *in vitro* (Fig S6A-B). In live cells, we observed a bimodal FRET distribution consistent with the existence of both unbound (80%, E = 0.40) and bound (20%, E = 0.83) populations (Fig 6B). In contrast, a duplex control showed only a single, low-FRET peak (Fig 6C; c.f. the *in vitro* data – Fig S6C). The absence of a high-FRET population for this construct, which is not a substrate for the polymerase, is consistent with the interpretation that the high-FRET population observed with the gapped-DNA construct is a result of bending induced by the endogenous full-length Pol.

Whilst the labelling scheme above discriminated well between the FRET signals arising from unbound and bound DNA, we could not resolve the smaller FRET difference between the binary complex and high-FRET ternary complex seen in our *in vitro* work (Fig S6B). We thus internalised the T(−18)T(+15) gapped substrate, which showed *in vitro* a larger FRET difference between the binary complex and the high-FRET ternary complex (Fig S6D-E). The resulting FRET histogram from live cells lacked a significant high-FRET peak, but did exhibit two low-FRET peaks, consistent with the presence of unbound DNA, and DNA in the binary complex (Fig S6F), and suggesting that little high-FRET ternary complex was present *in vivo*.

## Discussion

The combination of single-molecule FRET with both coarse-grained and all-atom molecular simulations has provided substantial mechanistic and structural insight into the recognition and binding of DNA substrates by Pol. We have characterised the structure and dynamics of multiple species present in solution: the substrate alone, the binary complex and the high-FRET ternary complex. Further, we have obtained evidence for the *in vivo* relevance of the bent binary complex, detecting its FRET signature in live cells.

We obtained a unique, solution-based, high-precision structure (RMSD = 3.8 Å) of Pol bound to a gapped-DNA substrate, containing upstream and downstream duplex DNA flanking a 1-nt gap (Fig 2B and 3A). Previous structural efforts lacked any downstream duplex DNA and so its position and the conformation of the substrate were unknown. Gapped DNA in the binary complex structure adopted a 120° bend (discussed further below).

The location of the upstream DNA in the docked structure agrees very well with existing co-crystal structures containing primer-template substrates. This supports our rigid-body docking approach, and the accuracy of our positioning of the downstream DNA on the fingers subdomain. This positioning conclusively rejects early propositions that the DNA might be channelled through the cleft formed by the fingers and thumb subdomains (33, 34). Our structure served as a starting point for all-atom MD simulations, which showed DNA dynamics in the binary complex, and identified transient DNA interactions with specific Pol residues. Some of these interactions involved residues implicated in previous biochemical studies, e.g. Y719 (17), providing a structural and mechanistic explanation for the experimental data; other residues (e.g. K549) revealed novel interactions that will merit further study.

Our docked structure showed that the downstream DNA was positioned very close to Y719 (Fig 3D), confirming its involvement in strand displacement. DNA Pol I shares a three-helix bundle (O, O1 and O2) structural motif with T7 RNA polymerase (35). This motif participates in DNA binding and strand separation (36), and includes conserved residues Y719, S717 and R789 in Bst (F771, S769 and R841 in *E. coli)*, which have been shown to be important for strand-displacement by Pol (17). This role for Y719 was further supported in our simulations, which showed the three-helix bundle (and particularly Y719) to be positioned between the template and non-template strands of the downstream DNA. The exact position of Y719 close to bases B(+1) and B(+2) on the downstream-template DNA is consistent with cross-linking data (37, 38).

We also identified residues that interacted with the downstream DNA (R779 and R784; Fig 3E). These residues are highly conserved, with published sequence alignments showing 29 and 48 out of 50 bacterial polymerase sequences containing a homologous residue at positions 779 and 784, respectively (37). The two residues are likely to be functionally complementary given their proximity in the structure and the similar interactions they form with downstream DNA in our simulations. Whereas our simulations indicate that R779 is more important for contacting DNA in the Bst Pol I, R784 may be the key residue in other bacterial polymerases that lack a positively charged residue at position 779, such as *E. coli* Pol. Interestingly, mutation of R784 to alanine (R836A in *E. coli)* has been shown to *increase* the binding of downstream DNA to the polymerase site (37, 39), possibly due to R784 contributing to the bending and distortion of downstream DNA, or reflecting an unfavourable orientation of the side chain in the DNA-Pol binary complex.

K549 is part of a conserved motif (K)<u>K</u>T present in 33 out of 50 bacterial polymerase sequences analysed (37). In our simulations, interactions with K549 appear to keep the template strand away from its non-template counterpart, which may facilitate strand separation. Radioactive competition assays and cross-linking experiments have shown that Pol forms contacts with the first 4 nucleotides of the downstream template strand (37), which are beyond the reach of the active-site residues (Y714, S717, Y719 and R789), but could be accounted for by interactions with K549. The identity of the amino acid(s) cross-linking to base +4 could not be identified in these studies, likely due to the dynamics of the template strand and the transiency of interactions with K549, both features being apparent in our simulations.

The binary complex structure from rigid-body docking suggested that the downstream DNA cannot be fully base-paired proximal to the Pol fingers (Fig 2D). This idea was supported by our MD simulations, in which 4-5 nt of the downstream DNA remained single-stranded for the majority of the simulation time (Fig 3B). Our quenchable FRET assay confirmed that the downstream DNA is indeed melted when bound by Pol (Fig 4). When carrying out Okazaki fragment processing or long-patch base excision repair, Pol must perform strand-displacement DNA synthesis, replacing the RNA primer / damaged DNA with newly polymerized DNA. Our data suggest that the strand-displacement process starts before any DNA synthesis, with up to seven nucleotides being melted upon Pol binding to the substrate.

Our *in vivo* single-molecule experiments unequivocally show that non-extendible gapped-DNA constructs are bent in live *E. coli*, unlike duplex DNA. The close agreement between the FRET signatures of the bent species in cells and *in vitro* suggests that bending is likely mediated by the endogenous full-length Pol binding, although the effect of other DNA binding proteins cannot be excluded. For both internalized labelled DNAs, we observed a higher proportion of the lowest FRET species (corresponding to unbound DNA) than expected from our *in vitro* binding data and the expected cellular Pol concentration (~400 nM (40)). The high abundance of the low-FRET molecules in cells may reflect the effect of intracellular conditions (e.g., the presence of free nucleotides that can transiently occupy the 1-nt gap), the involvement of other proteins that could compete with Pol for gapped-DNA binding, or a lower affinity of Pol for gapped substrates *in vivo*.

Previous *in vitro* studies observed the presence of two molecules of Pol bound to DNA substrates (41-43). We also observed Pol_2_-DNA species in our *in vitro* titrations (Figs S1 and S6), but not *in vivo*, suggesting that these complexes are unlikely to be important in the cellular context, where the presence of the 5’-nuclease domain in the full-length protein may inhibit dimer formation.

Gapped DNA in the binary complex structure exhibited a 120° bend (Fig 2B and 3A). DNA bending was also observed in the crystal structure of the mammalian gap-filling DNA polymerase β, where the ~90° bend observed was suggested to be important for the mechanisms of polymerisation and fidelity (6). Our data support the idea that bending may be a necessary mechanistic step for gap-filling polymerases, exposing more of the template base for interrogation by the incoming nucleotide. However, we propose bending may also play a role in substrate recognition and selectivity.

Our coarse-grained simulations on the free gapped DNA showed remarkable agreement with the smFRET data (Fig 5B) and have important implications for the binding mechanism of Pol. Since the breaking of the stacking interactions opposite the gap increases DNA bendability, unstacking will likely occur as a step on the path to Pol binding. In addition, the high flexibility of the unstacked DNA suggests that the substrate can adopt a close-to-final bent conformation even prior to Pol complex formation. The simulations also provide an explanation for Pol substrate specificity, specifically its increasing binding preference for gapped over nicked DNA, previously observed by gel shift assays (41) and ensemble anisotropy (44). This preference appears to arise from the increased flexibility of the gap over the nicked DNA, reflected in the different energy cost required for their bending. In this way, the substrate specificity is encoded in the structure and dynamics of the DNA substrate itself, allowing sequence-unspecific recognition of gapped DNA by Pol.

Interestingly, other forms of DNA modification can affect DNA flexibility; cytosine methylation reduces flexibility, while 5-formylcytosine (a substrate for base excision repair) was shown to increase flexibility (Ngo et al., 2016). Thus, it is likely that increased DNA flexibility may act as a general recognition signal for a variety of DNA repair processes.

Based on our results, we propose the following model for recognition and binding of a gapped DNA substrate by Pol involving conformational capture followed by an ‘on-protein’ rearrangement (Fig 7). The DNA substrate rapidly interconverts between stacked and unstacked states; the unstacked conformations are generally more bent and show increased fraying 1-2 nt around the gap. The Pol initially interacts with the upstream DNA while the substrate is in an unstacked state (conformational capture). This upstream region of the substrate resembles a primer-template structure, which is known to bind tightly to Pol (K_D_ < 1 nM; Turner, Grindley and Joyce, 2003) forming a sufficiently stable complex for crystallization (16, 18). This conformational selection step does not necessarily require the substrate to adopt the precise 120° bend angle seen in the binary complex; rather, the DNA conformational flexibility helps to avoid blocking binding through steric clashes. Having bound the upstream duplex, the downstream duplex is free to sample conformational space (as seen in the MD simulations on the binary complex; Fig 3A), docking to the protein, and fraying the additional 3-4 nts, resulting in the complete binding of the gapped DNA (K_D_ = 0.4 nM; Fig S1A). This proposed two-step binding mechanism comprises an initial conformational selection step in which the substrate is bound, followed by an ‘on-protein’ conformational search, in which the DNA and the protein both search conformational space

Other structure-specific DNA binding proteins which have been shown to interact with bent DNA (e.g. FEN1, Pol β) are also likely to exploit the conformational dynamics of their substrates for recognition and binding. Thus, the mechanism we propose of an initial conformational selection step, sensing the increased flexibility of the substrate DNA, followed by an ‘on-protein’ rearrangement, may be generally applicable to many structure-specific DNA binding enzymes. It is an attractive model for how these enzymes operate during DNA repair, where vast regions of undamaged DNA are searched rapidly to identify sites that need to be repaired to stop accumulation of toxic intermediates and mutations, and ensure normal cellular function.

**Figure 7.**
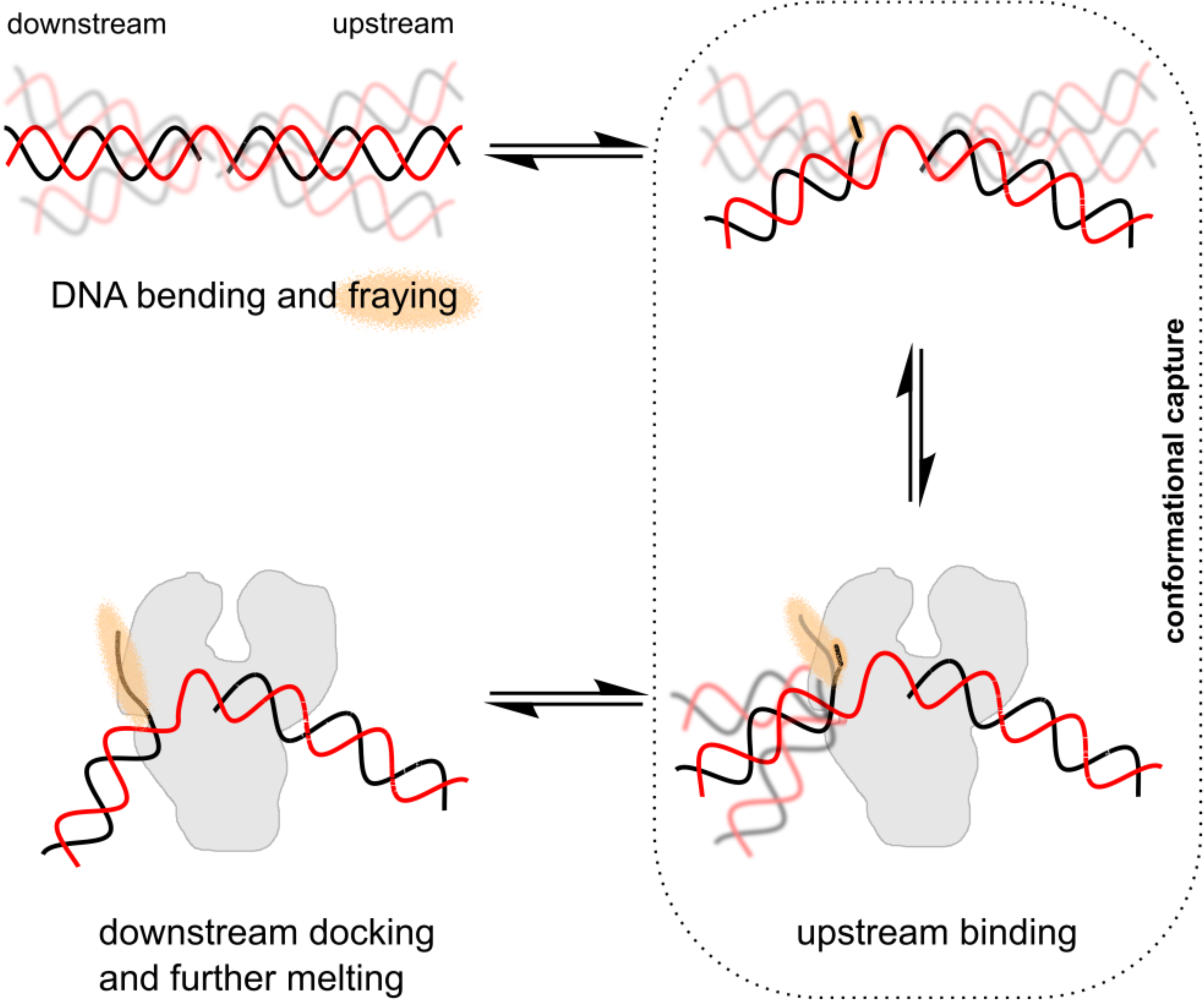
Gapped DNA recognition: conformational capture followed by an ‘on-protein’ rearrangement. Gapped DNA is dynamic adopting bent and frayed states (orange haze). Pol can bind to the upstream DNA when the downstream DNA conformation is not impeding the Pol (conformational capture of slightly bent states). Following binding of the upstream DNA, the downstream DNA now docks and is further melted, beginning the process of strand-displacement.

## AUTHOR CONTRIBUTIONS

Conceptualization, T.D.C., J.H. and A.N.K.; Software, T.D.C. and J.H.; Investigation, T.D.C., M.S., A.P., M.M., H.K., A.C. and J.H.; Resources J.P.K.D., P.C.B. and A.N.K.; Writing - Original Draft, T.D.C., M.S., and M.M.; Writing - Review and Editing, T.D.C., M.S., A.P., M.M., J.H., J.P.K.D., P.C.B. and A.N.K.; Supervision, T.D.C., J.P.K.D., P.C.B. and A.N.K. Funding acquisition, A.N.K.

## ACKNOWLEDGEMENTS

We thank Maria Musgaard for assistance with all-atom MD simulations, and are grateful for funding from: Lindemann Trust Fellowship (T.D.C.); Wellcome Trust (M.S.); German National Academic Foundation (Studienstiftung) and Phizackerley Senior Scholarship in Medical Sciences by Balliol College (A.P.); Marie Curie Career Integration Grant [#630992] (J.H.), UK EPSRC (A.P., M.M and J.P.K.D.); European Research Council (261227) and the UK BBSRC (BB/J00054X/1) (A.N.K.).

## METHODS

**Protein Expression, purification and labelling**. Pol variants were expressed from an N-terminal-His6, D424A construct and purified as described (10). The D424A mutation inhibits the proof-reading exonuclease activity. Briefly, a plasmid carrying the gene encoding Pol was transformed into HMS174 (DE3) cells, and single colonies inoculated in 25 ml LB, supplemented with 50 μg/ml carbenicillin. The cultures were grown overnight at 220 rpm and 37°C, and were used to inoculate 1 liter of LB supplemented with carbenicillin. The culture was grown to an ÜD600 of 0.6, at which point expression was induced with 0.5 mM Isopropyl β-D-1-thiogalactopyranoside (IPTG). After 2 hours of expression, the cells were harvested by centrifugation (20 min at 3000 rpm and 4 °C; GS-6R Beckman), resuspended in cold 50 mM Tris pH 7.5, and spun down in an ultracentrifuge (15 min at 8,000 rpm at 4 °C; Sigma 3K30, rotor 12150-H). Finally, the pellet was resuspended in lysis buffer (50 mM Tris pH 7.2, 300 mM NaCl, 1 mM β-mercaptoethanol, 10 mM imidazole, 2 mg/ml lysozyme and 0.02 mM Phenylmethane sulfonyl fluoride; PMSF). The cells were stored in the lysis buffer overnight at -80°C.

The frozen cells were thawed and fresh PMSF (25 μ!) was added. Cells were lysed by sonication (6 cycles of 5-sec ON and 10-sec OFF time) and the cell debris was spun down (20 min at 15,000 rpm at 4 °C; Sigma 3K30). The supernatant containing the cell lysate was combined with Ni-NTA resin (pre-equilibrated in buffer A: 50 mM Tris pH 7.2, 300 mM NaCl, 1 mM β-mercaptoethanol and 10 mM imidazole), and the protein allowed to batch-bind (1 hour at 4°C). The resin was spun down, resuspended in buffer A, applied to a plastic column and washed with buffer A containing increasing concentrations of imidazole (10, 20, and 27 mM). The protein was eluted in buffer A containing 100 mM imidazole, and the fractions analyzed by absorbance and SDS-PAGE. The concentrated fractions were pooled, and dialyzed into 50 mM Tris pH 7.2, 1 mM dithiothreitol (DTT) overnight at 4 °C. The dialyzed samples were combined in a 1:1 ratio with 2x glycerol storage buffer (80 % glycerol, 50 mM Tris pH 7.2, 2 mM DTT) and stored at -20°C.

Pol variants containing a single cysteine (C907+, C907S / K550C and C907S / L744C) were labelled using a maleimide derivative of Cy3B (GE Healthcare) as described (10). Briefly, purified Pol samples were reduced (5 mM DTT, 1 hr at 22°C), and DTT was removed by dialysis into 50 mM Tris pH 7.1, 0.12 mM tris(2-carboxyethyl)phosphine (TCEP). A two-fold excess of the Cy3B maleimide dissolved in DMSO was added to the protein sample and the reaction allowed to proceed overnight at 4 °C with gentle rocking. The reaction was quenched with 1 mM DTT, and applied to a heparin column (preequilibrated in heparin buffer containing 20 mM Tris pH 7.4, 1 mM ethylenediaminetetraacetic acid (EDTA), 2 % glycerol and 1 mM β-mercaptoethanol), washed with heparin buffer containing 50 mM NaCl, and the protein eluted in buffer containing 400 mM NaCl. Samples were dialyzed first into 1 liter of 50 mM Tris pH 7.4, 25 mM NaCl, 1 mM DTT, for 3x 1 hour, and then into 500 ml of the same buffer containing 40 % glycerol, overnight, before storing at -20 °C. Labelling efficiencies (typically ~80 %) were determined by UV-Vis absorbance, using extinction coefficients for Pol (58,790 M^−1^ cm^−1^ at 280 nm) and Cy3B (130,000 M^−1^ cm^−1^ at 570 nm), and taking into account the Cy3B absorbance at 280 nm.

**DNA labelling and annealing**. DNA oligonucleotides (oligos; Table S3) were prepared using automated synthesis (IBA GmbH), and labelled with NHS-ester derivatives of Cy3B (GE Healthcare) and Atto647N (Atto-tec) via dT-C6-amino linkers at selected positions according to the manufacturers’ protocols. Labelled oligos were purified by 20% polyacrylamide gel electrophoresis. Bands were visualized by UV-shadowing, cut, and extracted from the gel using an overnight crush and soak protocol at 4°C. The sample volume was reduced (by centrifugal evaporation) and buffer exchanged into TE buffer (Microbiospin6 columns, BioRad). Gapped-DNA substrates were assembled by annealing three single-stranded oligos (one from each group - DNA1, DNA2 and DNA3; Table S3) in annealing buffer, 20 mM Tris-HCl pH 8.0, 100 mM NaCl, and 1 mM EDTA. Samples were heated to 94°C and subsequently cooled to 4°C, in steps of 10°C over 45 min. Annealed substrates were stored at -20°C. For DNAs prepared for electroporation, the oligonucleotides were annealed in a low-salt annealing buffer (20 mM Tris-HCl (pH 8.0), 10 mM NaCl, 1 mM EDTA).

**Single-molecule FRET measurements**. Single-molecule FRET measurements were performed at room temperature using a home-built confocal microscope with 20 kHz alternating-laser excitation between a 532-nm (Samba, Cobolt, operated at 240 μW) and a 638-nm laser (Cube, Coherent, operated at 60 μW), coupled to a 60x, 1.35 numerical aperture (NA), UPLSAPO 60X0 objective (Olympus). For DNA-DNA measurements, labelled DNA was present at < 100 pM and unlabelled Pol (when present) at 3 nM concentration. For Pol-DNA measurements, both Pol and DNA were present at 100 pM concentration. Measurements were taken in ‘Pol buffer’, consisting of 40 mM 4-(2-hydroxyethyl)-1-piperazineethanesulfonic acid (HEPES)-NaOH, pH 7.3, 10 mM MgCl_2_, 1 mM DTT, 100μgml^−1^ bovine serum albumin, 5% (vol/vol) glycerol, 1 mM mercaptoethylamine. We recorded 3-6 datasets of 10 min for each distance measurement and combined for analysis. Photon streams in DD, DA and AA channels were recorded and processed using custom-written software (LabVIEW). Bursts were filtered for the correct labelling stoichiometry (45), and accurate FRET was calculated as described below. FRET histograms were fitted to single, double or triple Gaussian functions.

**Derivation of the multistate equilibrium model**. A three-species model, in which the observed low-, mid-, and high-FRET states correspond to DNA alone, DNA-Pol binary complex, and a dimer DNA-Pol_2_ respectively, could not account for the persistence of the mid-FRET signal at high Pol concentrations (Fig S1A). The simplest model that could account for all the data involved a second dimer species with a FRET efficiency indistinguishable from the DNA-Pol binary complex (Fig S1B). In this model, binding of Pol to the DNA, forming the binary complex (Pol:DNA*), is described by the association constant, K_1_=[Pol:DNA*]/[DNA]. Binding of a second Pol to the binary complex yields a mid-FRET dimer species (Pol_2_:DNA*) governed by the association constant *K*_2_=[ Pol_2_:DNA*]/[ Pol:DNA*]. This mid-FRET dimer can isomerize to the high-FRET dimer (Pol_2_:DNA**), a process described by the equilibrium constant *K*_3_=[ Pol_2_:DNA**]/[ Pol_2_:DNA*]. The corresponding dissociation constants are defined as *K*_D1_=1/*K*_1_ for the formation of the binary complex and *K*_D2_=1/(*K*_1_*K*_2_) for the formation of the mid-FRET dimer species.

The total DNA substrate concentration **is**:

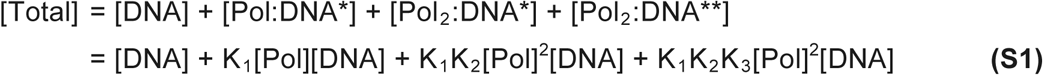

The fractions of each species as a function of Pol concentration are:

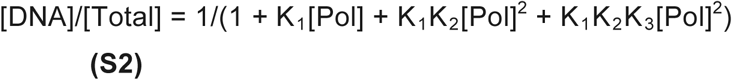

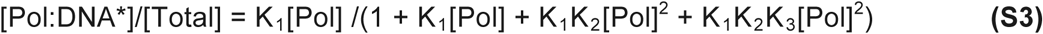

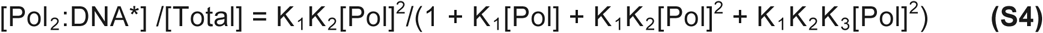

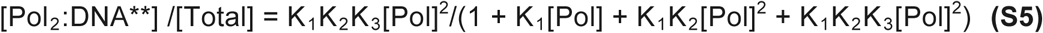

The fractional populations of the low-mid and high-FRET states are:

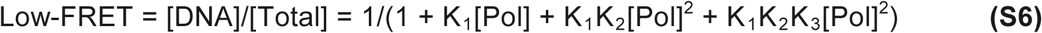

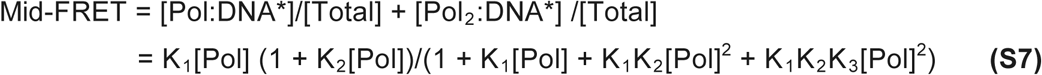

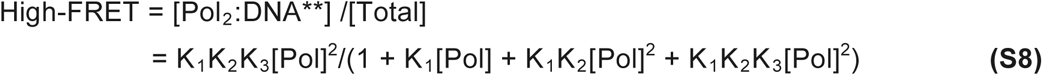

We used a global fitting approach to fit the variation in the fractional populations of the three FRET states simultaneously as a function of Pol concentration, and to determine the equilibrium constants *K*_1_, *K*_2_, and *K*_3_. From the equilibrium constants and their standard errors, we calculated the dissociation constants.

**Accurate FRET corrections**. The apparent FRET efficiency, E* was calculated from the DA and DD photon streams:

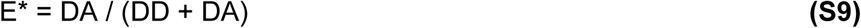

Similarly, the apparent stoichiometry, S* was calculated using:

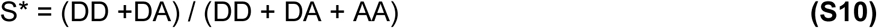

To obtain the accurate FRET efficiency, the raw photon streams were sequentially corrected for background counts, cross talk, and gamma / beta factors (which take into account the different detection efficiencies, quantum yields and excitation cross sections of the two dyes), as described (12, 13).

First, the three photon streams were corrected for background, which arises from impurities, Raman scattering from the solvent and dark counts in the detectors. For each burst, the corrected counts were calculated from the raw counts by subtracting the background count rate, multiplied by the length of the burst. Typical background count rates were 1-3 photons per ms.

After the background correction, the leakage fraction of the donor emission into the acceptor detection channel and the direct excitation of the acceptor by the donor-excitation laser were obtained. The correction factor for leakage (lk) was determined from the FRET efficiency of the donor-only population, E_don-only_:

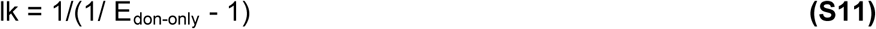

The correction factor for direct excitation (dir) was determined from the apparent stoichiometry value of the acceptor-only population, S_acc-only_:

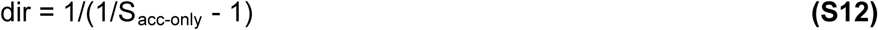

The DA intensities and the FRET efficiency and stoichiometry were then corrected as follows:

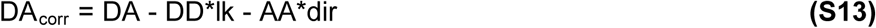

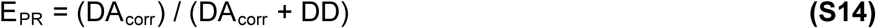

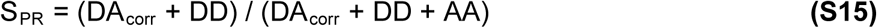

Finally, the gamma and beta parameters were obtained from a linear fit to a plot of 1/S_PR_ vs E_PR_:

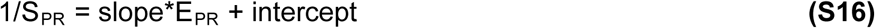

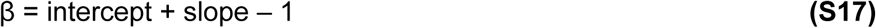

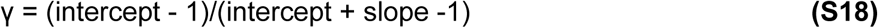

The fully-corrected accurate FRET efficiencies and stoichiometries, E and S are given by:

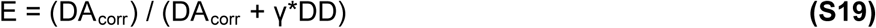

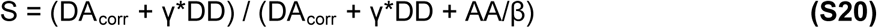

Accurate determination and application of all correction parameters was checked visually on the ES histograms, as all FRET populations should be located at S~0.5. Gamma and beta factors were determined separately for DNA-DNA and DNA-Pol measurements. This was necessary because of the significant difference in the quantum yield of the donor when attached to DNA or protein (see below and Table S4).

**Conversion of accurate FRET to distance**. Accurate FRET efficiency E, was converted to distance R, according to the equation:

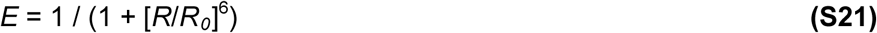

using experimentally determined values for the Förster radius, *R*_0_, which were calculated according to the equation:

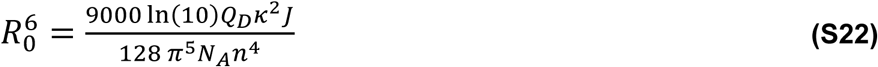

where *Q*_D_ is the quantum yield of the donor (which must be measured; see below), *N*_A_ is Avogadro’s number, and *n* is the refractive index of the medium. The term κ^2^ describes the relative orientation of the transition dipoles of the donor and acceptor. Its value lies in the range of 0-4, and it is often assumed to be equal to 2/3, which is the case when both fluorophores have unrestricted rotational freedom (46).The overlap integral *J* is a measure of the degree of overlap between the donor emission and acceptor excitation spectra (47), and can be calculated according to:

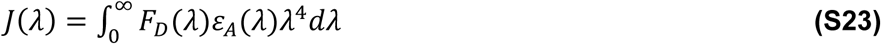

where *F*_D_ is the corrected donor fluorescence intensity at a particular wavelength λ, with the total intensity normalized to unity, and *ε*_Α_ is the extinction coefficient of the acceptor at the same wavelength.

Quantum yields were measured according to established methods (48, 49) for the following donor samples: free Cy3b-maleimide dye, Cy3b attached to gapped DNA (in the presence and absence of unlabelled Pol in a 1:1 molar ratio), and Cy3b attached to different positions of Pol (K550, L744, C907). Each sample was diluted from a glycerol stock to 5 μΜ final concentration in Pol buffer. Free Cy3b-maleimide dye was reduced with 10 mM DTT for 10 min prior to dilution. Absorbance at 490 nm was recorded for each sample, using a UV-visible spectrophotometer (Cary 50 Bio, Varian). An emission scan was taken of the same sample using a steady-state fluorimeter (PTI), exciting at 490 nm and recording at 510-700 nm. Samples were diluted and recordings repeated 5 times, to populate absorbance in the 0 to 0.1 region, where absorbance and emission are linearly related. The same procedure was applied to the reference dye, rhodamine 6G, dissolved in ethanol. The quantum yield of the donor dye was then calculated according to equation:

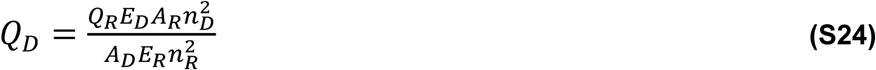

where *Q* is quantum yield, *E* is integrated emission across the whole spectrum, *A* is absorption at 490 nm, *n* is the refractive index of the medium, and *D* and *R* refer to the donor and the reference dyes, respectively. Established values were taken for the quantum yield of rhodamine 6G in ethanol (0.95; Magde, Wong and Seybold, 2002) and for the refractive index of ethanol (1.361). To calculate the overlap integrals, absorption spectra of the following acceptor samples were also measured: Atto647 free dye, Atto647N attached to DNA (in the presence and absence of Pol in a 1:1 ratio), and Atto647N attached to Pol. Samples were diluted in Pol buffer to 2 μΜ concentration, and absorption recorded at 400-710 nm. Both the absorption spectra of the acceptor, and the emission spectra of the donor (see above) were corrected for background, normalized, and the overlap integral calculated as in equation 3.3. The extinction coefficient of Atto647N at A_max_ was taken as 150,000 M^−1^cm^2^ (from manufacturer’s website; http://www.atto-tec.com). This allowed isotropic R0 values to be calculated (Equation S22), assuming orientational averaging (κ^2^ = 2/3) and the refractive index of water (n=1.333).

To test if orientational averaging is justified, steady-state anisotropies were measured (48). Samples were diluted to 100 nM in Pol buffer and excited with vertically polarized light at 532 nm (donor) or 638 nm (acceptor samples) in a steady-state fluorimeter (PTI). Fluorescence was measured through horizontally and perpendicularly oriented emission filters at 570 nm (donor) or 669 nm (acceptor samples) over 1 minute. Anisotropy values (Table S4) were calculated from the difference of vertically and horizontally polarized emission intensities, corrected for background and for the different sensitivities of the emission channel for vertically and horizontally polarized light.

**Distance calculations and accessible volume modelling of dye positions**. Accurate FRET efficiencies were converted to their corresponding distances using a FRET error of +/− 0.025 (determined from the standard error of the mean in three independent FRET measurements of the same sample), and using experimentally determined R_0_ values (Table S4). The *R*_0_ values used were 64.5 Å for DNA-DNA and for 59.0 Å for DNA-Pol distances, and the error in *R*0 was assumed to be the error that was propagated from the uncertainty in quantum yield determination of +/* 0.10. The experimentally determined distances correspond to FRET-averaged distances, *<R*_*DA*_>_*E*_ in the accessible volume model established by the Siedel laboratory (28). In this model, dye rotation occurs faster than the FRET process, but the position of the dye is fixed on the timescale of FRET. Other dye modelling methods based on Bayesian statistics can also be used (Muschielok et al., 2008).

The FRET-averaged distance is calculated by averaging the distances between individual dye positions in the donor and acceptor accessible volumes. For rigid body docking, these FRET-averaged distances were converted to distances between mean dye positions *R*_*mp*_, using a third-order polynomial function:

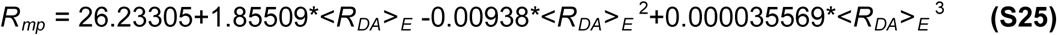

that was established by calculating *R*_*mp*_ and *<R_*DA*_>*_*E*_ values for pairs of dyes at different positions along a double-stranded DNA using, as described (29). The radii, linker lengths and linker widths of Cy3B and Atto647N dyes were estimated from their structures in silico using ChemDraw (Perkin Elmer) and are given here (in Å):

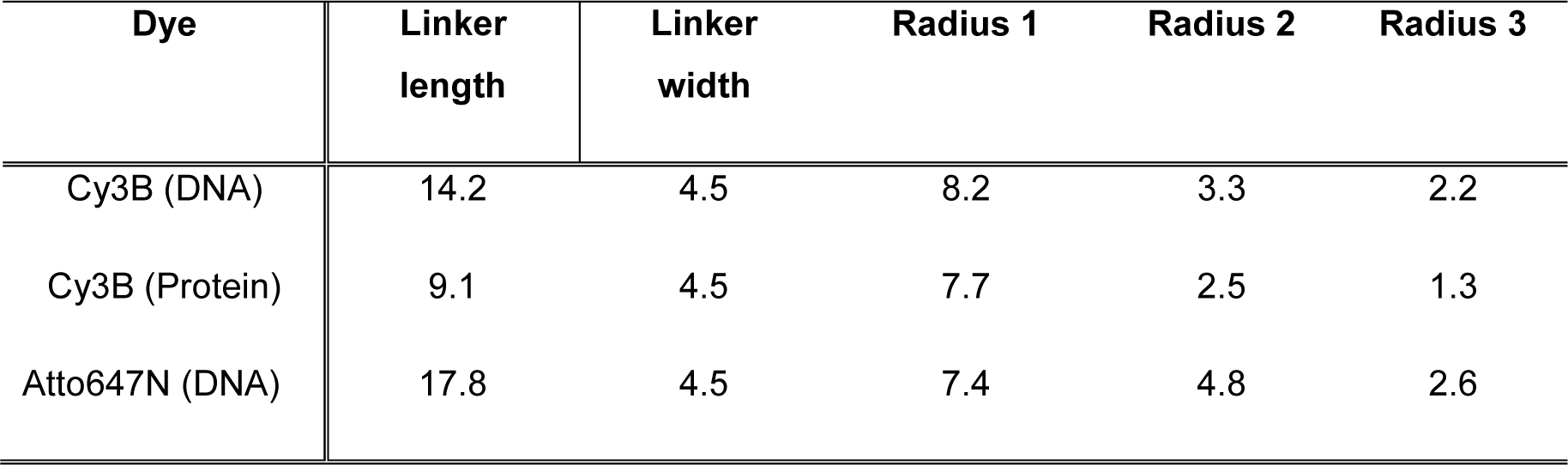

We used the accessible volume (AV) algorithm of the FPS software (29) to model the mean positions of the dyes for each Pol and DNA attachment site. The attachment points were taken to be the S atoms of Cys residues, and the C7 atoms of dTTP residues. For quFRET (see below) we calculated the percentage of the donor accessible volume that overlapped with the acceptor accessible volume using custom written code in MATLAB (Mathworks). Accessible volume elements were counted as overlapping if the distance between them was smaller than the lattice spacing used to calculate the initial AV.

**Rigid body docking**. The polymerase structure was obtained from the *Bst* X-ray crystal structure (PDB code 1L3U; Johnson *et al*., 2003). DNA was removed and Cys substitutions were introduced at positions K498, V692 and A855 (corresponding to *E. coli* residues K550, L744 and C907) using the PyMOL Molecular Graphics System, Version 1.8 Schrödinger, LLC. B-DNA models of the upstream and downstream DNA were made using 3D-DART modelling server (51) and were truncated at the gap-proximal ends by 3 base-pairs each for the purposes of rigid-body docking. Three-body rigid-body docking with Pol, upstream and downstream DNA structures was performed in the FPS software (29) using the calculated *R*_*mp*_ distances. Docking was repeated 1000 times from different starting configurations of the binding partners, using a clash tolerance of 6 Å. This treatment generated several clusters of structures, which were distinguished by the different RMSD values relative to each other, and the different goodness of fit to experimental data assessed by the reduced chi-squared parameter (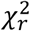). Structures with 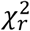; values above 6 were rejected, and one structure from each of the remaining clusters was further refined using a clash tolerance of 2 Å, and then again using a tolerance of 1 Å, during which steps the AV clouds were recalculated. The structure with the lowest 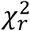; was taken, and the *R*_*mp*_ distances from the model back-converted to <*R*_*da*_>_*e*_ and FRET efficiency values, to compare with the experimental FRET data. For precision estimation, 100 bootstrapped structures were generated from the best model, using a clash tolerance of 1 Å. The coordinates of each P atom were extracted, and the RMSD of each P atom calculated across the 100 bootstrapped structures.

To compare the position of the upstream DNA in the docked structure with the crystal structure, the protein components of the FRET-restrained and crystal structures were aligned in PyMol. The RMSD of the upstream DNA fragment between the two structures was calculated as for bootstrapped structures, but across all P atoms. The DNA structure in complex with the Pol dimer was obtained using the same procedure as the Pol-DNA structure, but with no polymerase present in the docking. In the case of the DNA structure in the absence of Pol, the DNA fragments were at their full length, and with an additional distance restraint of 5 +/− 2.5 Å imposed between the C atoms in the template strand opposite the gap, to account for the covalent link between the two.

**All-atom molecular dynamics simulation - Model preparation**. The protein atoms and the catalytic magnesium ion were extracted from the *Bst* X-ray structure PDB file (code 4BDP; Kiefer et al. 1998). The online server ‘WHAT IF’ was used to check for errors in the PDB, and build the missing side chains into the structure (52, 53). PyMol was used to align the FRET-restrained structure with the X-ray structure, based on the protein component only. The downstream DNA in the docked structure was extended to its full length, and its template strand linked to the template strand of a 5-nucleotide fragment of upstream DNA from the X-ray structure (which also includes the templating nucleotide and one nucleotide downstream). Basic molecular sculpting was performed such that the conformation of the DNA backbone was not significantly disturbed, and no steric clashes occurred with the polymerase, which resulted in 6 base-pairs of downstream DNA being unpaired. The upstream DNA fragment was then extended using the sequence of upstream DNA from the docked structure. This step was justified by the excellent agreement in the position of the upstream DNA between the X-ray and docked structures (Fig S2B). For DNA-only simulations, DNA models were generated using the 3D-DART server (51). DNA atoms were extracted from the PDBs and terminal phosphate groups removed using PyMol. In the case of gapped DNA, the central nucleotide was removed and a 5’ phosphate group generated instead.

**Force fields and parameters**. All complex simulations and high-temperature DNA simulations were run using Amber ff99sb force field (54) with modified nucleic acid parameters (parmbsc0; (55, 56). The crystal structure control and the DNA-only simulations, were run using Amber ff99sb-ILDN with Amber94 nucleic acid parameters (57). No parameters were available in either force field for the 5’ phosphate groups of DNA, as these are usually missing in crystal structures due to their high flexibility. The phosphate group had to be modelled at the 5’ end of the gap in our DNA substrate, as it is both physiologically relevant and present in our single-molecule experiments. Therefore, the force fields were modified by assuming that the parameters of the β-phosphate of free ADP available online (http://research.bmh.manchester.ac.uk/bryce/amber/), are a reasonable approximation for the α-phosphate of the gap-proximal dDTP.

**Simulation conditions**. All simulations were carried out using Gromacs 4.6 (58). The X-ray structure control simulations and complex simulations were done using explicit solvent (TIP3P) in a triclinic box, with a minimum 10-Å solvent edge, in the presence of 10 mM MgCl_2_. The system was neutralized with addition of magnesium ions, and energy-minimized using steepest descent minimization. In order to stabilize the temperature of the system, equilibration was performed in the NVT ensemble for 100 ps, with the temperature of 298 K maintained using a Berendsen thermostat (59). Next, the pressure of the system was stabilized by equilibration in the NPT ensemble for 1 ns, with the temperature of 298 K and the pressure of 1 bar retained using a V-rescale thermostat (60) and a Berendsen barostat (59), respectively. During equilibration, DNA, protein heavy atoms and the catalytic magnesium ion were position-restrained with a force constant of 1,000 kJmol^−1^nm^−2^. DNA was equilibrated for an additional 10 ns with protein heavy atoms restrained, under the NPT conditions. Atom velocities were preserved between the equilibration steps, and between equilibration and production steps. Unrestrained production was finally allowed to run for 100 ns, with the temperature of 298 K and the pressure of 1 bar maintained by the V-rescale thermostat and a Parrinello-Rahman barostat (61). Periodic boundary conditions and the Verlet cutoff scheme were used, and long-range electrostatic interactions were accounted for by the Particle-Mesh Ewald method (62). All bonds were treated as constraints with the LINCS algorithm, resulting in a time step of 2 fs. Coordinates were saved to an output trajectory every 5 ps. Repeat simulations were carried out using different randomly numbered seeds, generating different initial atom velocities each time.

In the case of full-length DNA-only simulations, the conditions were the same except that a square box was used, with dimensions equal to the length of the DNA plus a 10-Â solvent edge. The NVT and NPT equilibration steps were performed, and the production times were 20 ns. In the case of high-temperature DNA simulations carried out as part of model preparation, the conditions were the same as for the complex simulations except that the temperature during the equilibration and production runs was 400 K, and the production times were 2 ns. All DNA heavy atoms were position-restrained during the production runs, except for the 6 base pairs in the protein-proximal, downstream part of the DNA, which were unpaired in the starting configuration.

**Analysis**. All analysis was carried out using Gromacs 4.6 or 5.0, and VMD (63). Trajectories were repaired for periodic boundary conditions, and processed to include only every 10th frame, corresponding to 50-ps steps. Maps of occupancy of DNA and of polymerase residues during the simulation were created with VMD’s volmap density function, using an isovalue of 0.001. RMSD and end-to-end distance measurements were done using standard functions in Gromacs. The flap-to-template H-bonds were quantified by measuring the number of bonds at any one time in the simulation, using a distance cut-off of 0.33 nm, and an angle cut-off of 30°. The position of residue Y719 relative to the DNA was calculated by measuring the distance between the centers of mass of Y719 side chain and individual DNA residue base moieties. Pol-DNA interactions were detected by measuring the minimum distance between any nitrogen atom of a specific Pol residue and a specific phosphorous atom in DNA, during the entire simulation. Distances below 0.4 nm were taken as indicating an interaction.

**Coarse-grained molecular dynamics simulation of DNA substrates using oxDNA**. DNA substrate systems were simulated using oxDNA, a nucleotide-level coarse-grained model of DNA in which each nucleotide is modelled as a rigid body. The oxDNA model has been described in detail (24, 64) and is implemented in a simulation package which is available for download (http://dna.physics.ox.ac.uk/). It was designed to reproduce the thermodynamic and mechanical properties of both single- and double-stranded DNA (24, 25), and has proven powerful in predicting the kinetics of the basic dynamical processes in DNA systems (27, 65, 66). Therefore, it is particularly suited for probing the structure and dynamics of the DNA substrates in this study. The oxDNA interaction potential consists of terms representing the backbone connectivity (modelled as a finitely-extensible nonlinear elastic spring), excluded volume, hydrogen bonding between Watson-Crick (WC) complementary base pairs, stacking between adjacent bases along the chain, coaxial stacking between non-adjacent bases, and cross stacking (Fig S5E). Aside from backbone connectivity and excluded volume, all interactions are anisotropic, depending on the relative orientation of the nucleotides. Orientational modulations of the stacking potential favors the bases to form coplanar stacks, and hydrogen bonding can occur between complementary WC base pairs when they are anti-aligned, leading to the formation of double-helical structures for which the helical twist arises from the different length scales of the backbone separation and the optimal stacking separation. Within oxDNA, the bases in the single-stranded DNA can stack/unstack and the strengths of hydrogen bonding and stacking interactions depend on the identities of the interacting bases (64). It has been parameterized for a NaCl concentration of 0.5 M, similar to the experimental buffer conditions in this study.

We performed 100 simulations of 10^8^ steps each for each of the gapped, nicked and duplex DNAs, with interaction energies and configurations sampled every 10^3^ steps. The time step was 0.005 simulation units, where one simulation unit implies a time of 3.03 ×10^−12^ s. The temperature was set to 295 K, and an Andersen-like thermostat was used (67). Particle velocities were refreshed every 10^3^ steps from the Maxwell distribution corresponding to the simulation temperature, with fixed probabilities of 0.02 and 0.0067 for the linear and angular velocities, respectively.

The bend angle was calculated from the vectors placed along the midlines of the two helical segments, as described previously (68). The relative free energies were calculated from the MD trajectories, as follows:

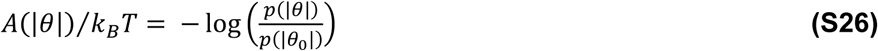

where *A*(|*θ*|) is the free energy, *k*_*B*_ is the Boltzmann constant, *p(|θ|)* is the observed probability density for the DNA adopting a bend angle |*θ*|, and | *θ*_0_| is the reference bend angle, for which *A*(|*θ*_0_|) = 0.

FRET efficiencies were calculated from the molecular dynamics trajectories, by adapting the accessible volume (AV) model for dye positions detailed above. Briefly, a grid of points was produced around the DNA base attached to the dye, with the spacing between grid points set to half the smallest dye dimension (see table above). Points were excluded if their distance to a base or backbone site, was smaller than the sum of the dye radius and the excluded volume radius of the base or backbone, respectively. This overlap check was repeated with the three different dye radii and the resulting AV clouds were combined (Fig S5G). A position that could accommodate all three dye radii was therefore weighted three times more than a position that could only accommodate one. The FRET efficiencies were averaged over all dye distances for each configuration and then again over all configurations in our molecular dynamics trajectories (of length ~15μs).

**Single-molecule FRET measurements of DNAs in living bacteria**. Gapped and duplex DNAs were internalized into electro-competent DH5α *E. coli* cells (Invitrogen) using electroporation (30). Cells were diluted 1:1 with sterile milli-Q water and stored at -80°C. For each electroporation experiment, 20 μL of electrocompetent cells were used. DNAs were stored in 2μΜ stocks in low-salt annealing buffer at -20 °C. For each experiment 0.25 μL of DNA and 0.2 μL of 50 mM EDTA were added to 20 μL electrocompetent cells and incubated on ice. The mixture of electro-competent cells and labeled DNAs was transferred into a pre-chilled electroporation cuvette (0.1 cm gap cuvette, Bio-Rad) and placed into an electroporator (MicroPulser, Bio-Rad). An electric field of 1.4 kV/cm was applied for electroporation. About 500 μL of super optimal broth with catabolite repression (SOC) was added immediately after electroporation. Cells were recovered for 3 min at 37°C. After recovery, cells were harvested by centrifugation at 3300 g for 1 min at 4°C and washed 5 times with 500 μL phosphate buffered saline (PBS). Cells were resuspended in 150 μL PBS and placed on 1% agarose pads before imaging. The agarose pads were made from ~300 μL of M9 medium containing 1% (v:w) BioRad Certified Molecular Biology Agarose on a coverslip. About 3 μL of cells were pipetted onto the agarose pad, and another coverslip was added on top. The slide/agar/slide sandwich was inverted and placed on the microscope with the side containing the cells closest to the objective.

Live-cell imaging was performed on a customized inverted Olympus IX-71 microscope equipped with a 532 nm DPSS laser (MGL_III-532-100mW, CNI). Laser light was collected into a single-mode optical fiber (Thorlabs, Newton, NJ, USA) and collimated before focusing on the objective. Cells were imaged using highly inclined thin illumination (HILO, Tokunaga et al., 2008) by adjusting the position of the focused excitation light on the back focal plane of the objective. Cellular fluorescence was collected through the same objective, filtered to remove excitation light through a long-pass filter (HQ545LP, Chroma) and a notch filter (NF02-633S, Semrock), and spectrally separated by a dichroic mirror (630DRLP, Omega). Donor and FRET channels were imaged onto separate halves of the chip of an electron-multiplying charge-coupled device camera (iXon+, BI-887, Andor). The illumination for brightfield images comprised a white-light lamp (IX2-ILL100, Olympus) and condenser (IX2-LWUCD, Olympus) attached to the microscope. Movies and images were recorded using manufacturer’s software. Measurements were performed in green continuous-wave mode using an excitation power density of 38 W/cm^2^ and 20 ms exposure time.

Custom-written MATLAB software was used to analyze single-molecule tracking and diffusion in live *E. coli* as previously described (30, 40). Briefly, the PSFs in donor and FRET channels in each movie frame were fitted by a 2D elliptical Gaussian (free fit parameters: x/y position, x/y width, elliptical rotation angle, amplitude, background) using initial position guesses from applying a fixed localization-intensity threshold on the bandpass-filtered fluorescence image (69). Tracking was performed in the FRET channel by adapting the MATLAB script based on a published algorithm (70). Localized PSFs were linked to a trajectory if they appeared in consecutive frames within a window of 7 pixels (~ 0.69 μm). This window size ensures 98% of steps are correctly linked for an apparent diffusion coefficient of 1.0 μm^2^/s at 20 ms exposure time. To account for transient disappearance of the PSF within a trajectory due to blinking or missed localization, we used a memory parameter of 1 frame. To eliminate noise, only molecules appearing in 5 consecutive frames were included in the analysis. The donor channel was mapped onto the FRET channel using a transformation matrix. FRET values, E*, were obtained from co-localized PSFs by calculating the ratio of photon counts in the FRET channel over the sum of photon counts in both channels for each single-molecule (71); E* = pc_FRET_/( pc_FRET_ +pc_Donor_), pc_FRET/Donor_: photon counts in FRET and donor channel, respectively.

Fluorescence overlay images were obtained by overlaying the donor and FRET fluorescence channels colored green and red, respectively. The green fluorescence channel was transformed onto the red fluorescence channel. All transformation matrices were based on a calibration matrix generated each day where fluorescent beads were mapped from the green onto the red fluorescence channel.

**Quenchable FRET (quFRET) experiments**. quFRET experiments were performed as per our standard smFRET confocal experiments (see above). DD, DA and AA photon streams were recorded and used to calculate the uncorrected FRET efficiencies (*E\** - equation S9) and stoichiometries *(S\** - equation S10) of filtered bursts, which correspond to individual molecules. These values were plotted as a two-dimensional histogram, and the number of bursts in the mid-S (0.4 < S* < 0.8) and low-S (S* < 0.4) regimes counted. One dimensional histograms of *E\** were produced from projections of the mid-S data onto the *E\** axis. The quFRET assay offers two related readouts for DNA melting: an increase in the absolute number of mid-S bursts, and an increase in the relative proportion of mid-S bursts compared with low-S bursts; the latter is a more robust measure, being independent of sample concentration and measurement time.

